# Global Distribution of Coronaviruses among Bat Populations detected using Molecular techniques, A Systematic Review

**DOI:** 10.1101/2025.03.01.640975

**Authors:** John A. Mulemena, Patson Sichamba, Walter Muleya, Benjamin Mubemba, Simbarashe Chitanga, Edgar Simulundu, Katendi Changula

## Abstract

Surveillance of bat coronaviruses (CoVs) is of public health importance, as accumulating evidence suggests that bats are hosts of the three significant pandemic viruses, namely Severe Acute Respiratory Syndrome Coronavirus (SARS-CoV), Middle East Respiratory Syndrome Coronavirus (MERS-CoV), and SARS-CoV-2. Studies focused on identifying different species of bat CoVs may have information cardinal for effective prevention and control of emerging zoonotic diseases. We conducted a systematic review using selected keywords (Surveillance, detection, identification, discovery, isolation, characterization, molecular methods, and Bat coronaviruses) to evaluate molecular studies on CoVs in bats. A total of 790 articles were found using the advanced search strategy of the PubMed database. Following the selection criteria, a total of 127 articles were finally chosen for full-text evaluation. Out of the total of 54 countries examined, China emerged as the country with the highest number of studies, accounting for 26% (n = 33). The sample categories consisted of faecal, urine, guano, blood, tissues, oral, and rectal swabs. The molecular detection approaches included polymerase chain reaction (PCR)-based techniques using species-specific, genus-specific, or broad-range primers. Approximately 94.5% (n = 120) of studies used PCR assays that amplified the partial RdRp gene of length ranging from 123 to 440 bp, followed by amplicon sequencing using either Sanger or next-generation sequencing technologies. Full genome sequencing was only performed in approximately 33.9% (n = 43), with metagenomics approaches being used in 15.7% (n = 20) of the studies. The higher positivity rate of bat CoVs were detected in Asia. Globally, the most predominant bat species which tested positive for CoVs were *Rhinolophus*, *Myotis*, *Miniopterus*, *Scotophilus*, *Eidolon*, *Chaerephon*, *Hipposideros*, and *Desmodus*. Continuous bat coronavirus surveillance using molecular methods and full genome sequencing is of utmost importance in detecting and characterizing viruses at molecular level and establishing the genetic diversity of new and circulating viruses.

## 1.0 Introduction

Bats are a taxonomically diverse assemblage of mammals that are categorized within the taxonomic order *Chiroptera.* Moreover, bats exhibit a broad geographic distribution, and substantial population sizes, ranking them as the second most diverse group of mammals after rodents (1). Bats have the capacity to serve as reservoir hosts for a variety of pathogens due to the diverse cell tropism they exhibit, their remarkable longevity, immunity to a wide range of viruses that afflict humans, and the heightened potential for viral transmission resulting from increased interaction between bats and humans (2–6). Several zoonotic viruses have originated from bats including paramyxoviruses, filoviruses, lyssaviruses, Hendra viruses, Nipah viruses, Ebola viruses and CoVs (7), (8). The initial global outbreak of SARS-CoV in 2003, widely believed to have originated from bats, led to an increased focus on CoVs’ surveillance in bat populations (9). The first identification of bat CoVs occurred in China in 2005 (10). Coronaviruses are a type of enveloped virus characterized by a positive-sense single-stranded RNA molecule, ranging in size from 27-32 kb. They are classified within the *Coronaviridae* family, which encompasses approximately 54 distinct species (11). The genome of CoVs encodes four structural proteins, namely spike (S) glycoproteins, envelope (E), membrane (M), and nucleocapsid (N), along with non-structural and accessory proteins (12). Most CoVs exhibit a broad host range across diverse animal species and possess the ability to cause a spectrum of diseases affecting the respiratory, gastrointestinal, hepatic, and neurological systems (13–16) (17). The grouping of these viruses is determined by their shared characteristics in terms of their replicase genes and the complement of accessory genes they possess (18). Hence, the viruses are categorized into four distinct genera, namely *Alphacoronavirus, Betacoronavirus, Gammacoronavirus,* and *Deltacoronavirus* (11). In general, it has been observed that *Alphacoronavirus* and *Betacoronavirus* primarily infect mammalian hosts, whereas *Gammacoronavirus* and *Deltacoronavirus* infect avian hosts (19). A common naming convention classifies detected bat CoVs as related to the known human CoVs. Thus, numerous studies have detected CoVs identical or closely related to SARS-CoV-1, (20), MERS-CoV (21), (22) and SARS-CoV-2 (23), (24).

Globally, pathogens have been detected using a variety of detection and surveillance methods, including molecular approaches (genomic surveillance), which are based on nucleic acid amplification and sequencing. In CoVs, the targeted sequences include genes coding for structural proteins such as spike (S) glycoproteins, envelope (E), membrane (M), nucleocapsid (N), and non-structural genes such as RNA-dependent RNA polymerase (RdRp), Helicase (Hel), ORF1a and ORF1b (25).

Despite the growing body of research on CoVs in bat populations, our understanding of their global distribution, diversity, and the methodologies used for their detection remains incomplete and fragmented. Current studies are often limited to specific regions and utilize varying detection techniques, leading to inconsistencies and gaps in the data. Furthermore, the main motivation behind nearly all viral surveillance conducted in wildlife is frequently prompted by disease outbreaks, resulting in a limited number of studies centred on wildlife and hindering the assessment of ecological dynamics of the bat CoVs genetic diversity, and patterns of zoonotic disease transmission (26).

This article provides a comprehensive analysis of surveillance studies that have utilized molecular techniques for the identification of bat CoVs globally. It explores the detection pipeline in the context of genetic analysis used for bat CoVs, the types of specimens analyzed, and the global distribution of bat species along with their associated CoV genera.

## 2.0 Methodology

### 2.1 Literature Search Strategy

This study adheres to the guidelines outlined in the Preferred Reporting Items for Systematic Reviews and Meta-Analysis (PRISMA) statement (27). The articles were sourced from the PubMed database (https://pubmed.ncbi.nlm.nih.gov/). The literature search was last conducted on 6^th^ December 2022 using selected keywords and Boolean connectors such as “surveillance,” “detection,” “identification,” “discovery,” “isolation,” “characterization,” and “molecular methods,” and “Bat coronaviruses.”

### 2.2 Study Selection

For the analysis, we exclusively considered primary publications written in English that specifically addressed the detection of bat coronaviruses. We also considered only publications that provided detailed information on the types of samples used, the specific molecular techniques employed for screening and detecting viral RNA, as well as the sequencing platform utilized. The initial screening of the articles involved an assessment of their title and abstracts. Subsequently, a comprehensive evaluation of the full text was conducted to determine their inclusion or exclusion, in accordance with predetermined criteria for inclusion and exclusion.

### 2.3 Inclusion and Exclusion Criteria

The study included original peer-reviewed articles in the English language, which used molecular methods to detect coronaviruses in bats. The exclusion criteria were articles focusing on human samples, reviews, opinion articles, duplicated articles, and conference abstracts.

### 2.4 Data Extraction and Data Items

Data were collected from articles using custom data extraction forms developed using Excel to capture key study variables like first author’s surname, article year of publication, study design, year of sample collection, nature of the specimen collected, molecular detection methods used, bat species found to contain the coronavirus, the total number of samples, number of positive samples, genus type, and country of study. The recorded variables were counter-checked by an independent researcher to avoid biases. The extracted data were securely kept in Google Drive, with limited access granted just to the research team. Periodic data backups were conducted to mitigate the risk of data loss.

### 2.5 Quality Assessment

The selected articles were subjected to quality assessment as designed by Joanns Briggs Institute (28) McMaster Critical Review Form (29). An independent reviewer assessed the outcomes of the search autonomously. Only articles, which met the criteria, were selected for final analysis.

### 2.6 Statistical Analysis

Data were analysed using Microsoft Office Excel 2018 and Stata software IC 15.0. Descriptive statistics such as frequency and percentages were performed.

## 3.0 Results

### 3.1 Study Selection and Characteristics

A total of 790 articles were retrieved using the advanced search strategy of PubMed followed by an assessment for eligibility. A total of 590 articles were excluded after being screened by title and abstract due to a lack of information on molecular diagnosis. The remaining 200 articles were then subjected to full-text evaluation, 127 articles were ultimately chosen and included, with 73 articles excluded because they did not focus on the detection of CoVs in bats, while some were review articles (Figure 1). Notably, 85 out of 127 (67%) of the articles were published between 2016 and 2022 (Figure 2).These studies were from 54 countries with China making up 26 % (33/127) of all studies (Figure 3).

**Figure 1:**
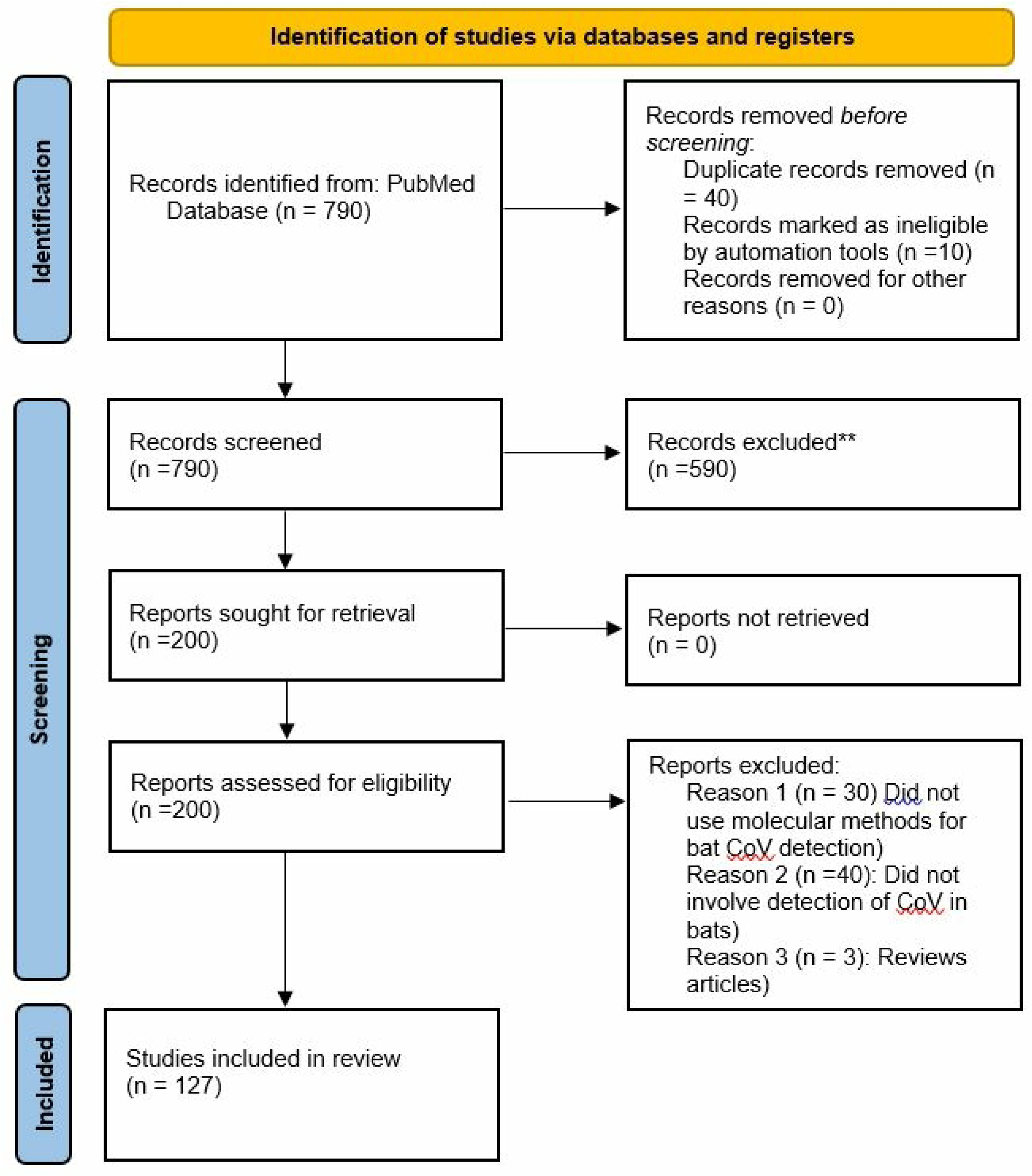
Flow diagram for the study selection process of publications included in the analysis.

**Figure 2:**
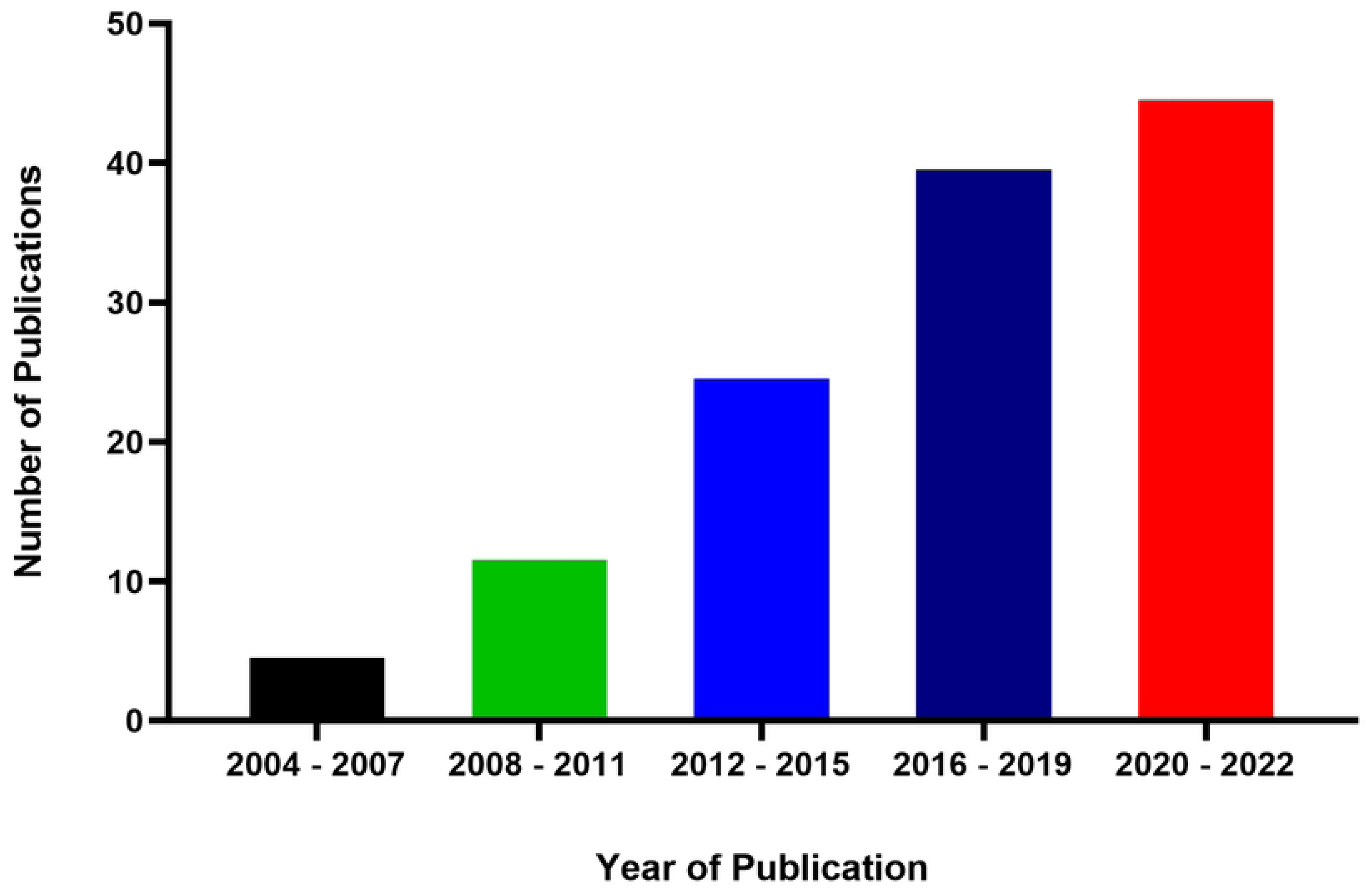
A bar graph showing the number of publications against the years the publications were published.

**Figure 3:**
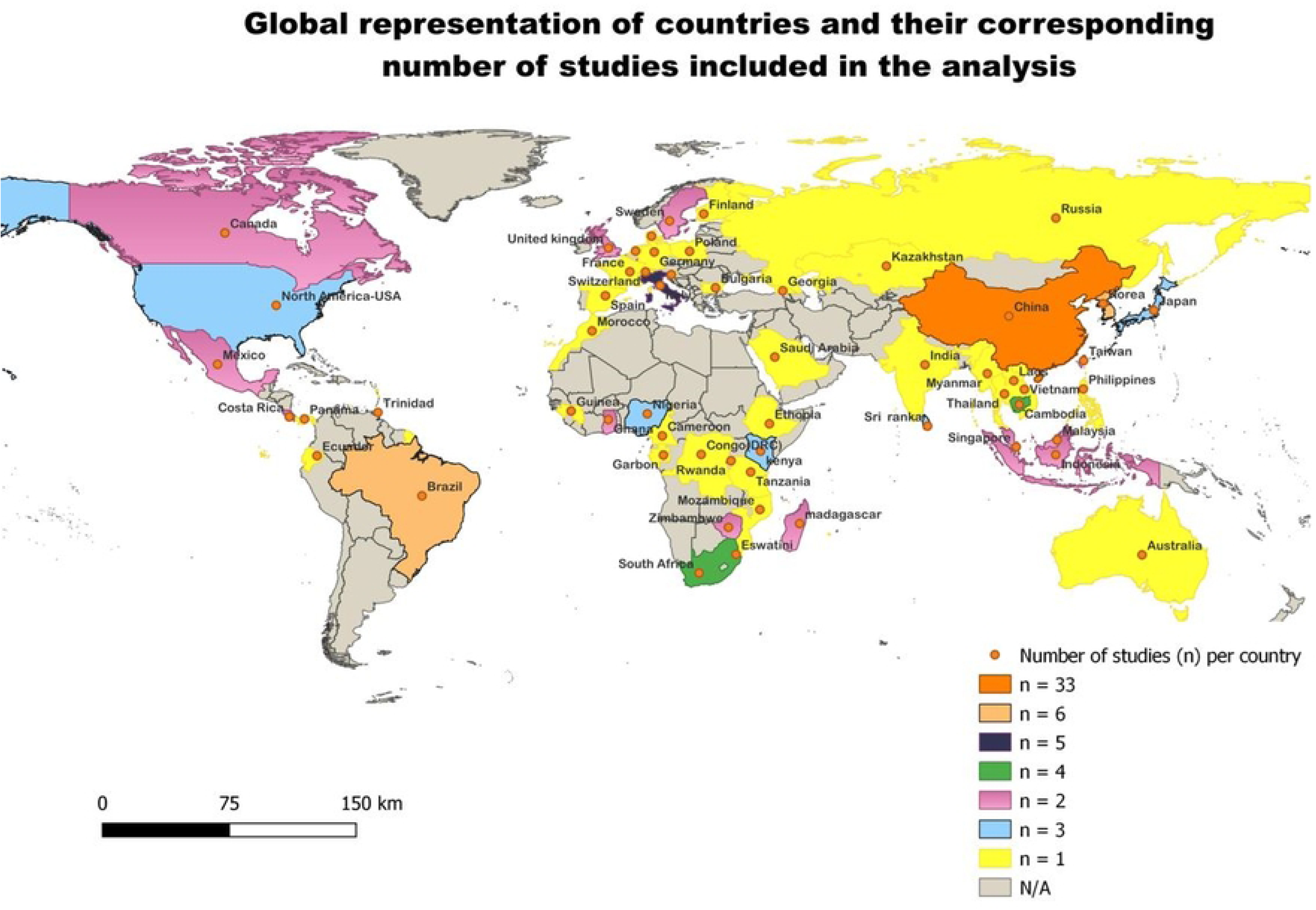
Global representation of selected countries included in the analysis.

The reviewed studies, published between 2005 and 2022 (Figure 2), reflect the growing focus on bat coronavirus surveillance. This increase has been driven by the emergence of three respiratory zoonotic diseases.

### 3.2 Sampling strategy

The sampling strategies used were non-invasive and/or invasive sampling techniques. The majority of studies used non-invasive sampling techniques (Table 1). Non-invasive sampling techniques involved collection of oral and rectal swabs, faecal material, and urine. (Figure 5). Invasive sampling techniques involved sacrifice of the bats and collection of blood and bat tissues. The majority of studies incorporated multiple sample types, with faecal samples being the most frequently collected sample, accounting for approximately 61.4% of the studies (Figure 5). The utilized specimen categories encompass guano, faeces, urine, nasal secretions, throat swabs, and saliva or oral swabs, which are considered non-invasive. Likewise, liver, intestines, brain, lungs, spleen, kidneys, and heart specimens are employed, which are classified as invasive. About three-quarters of the studies used non-invasive sampling.

**Figure 4:**
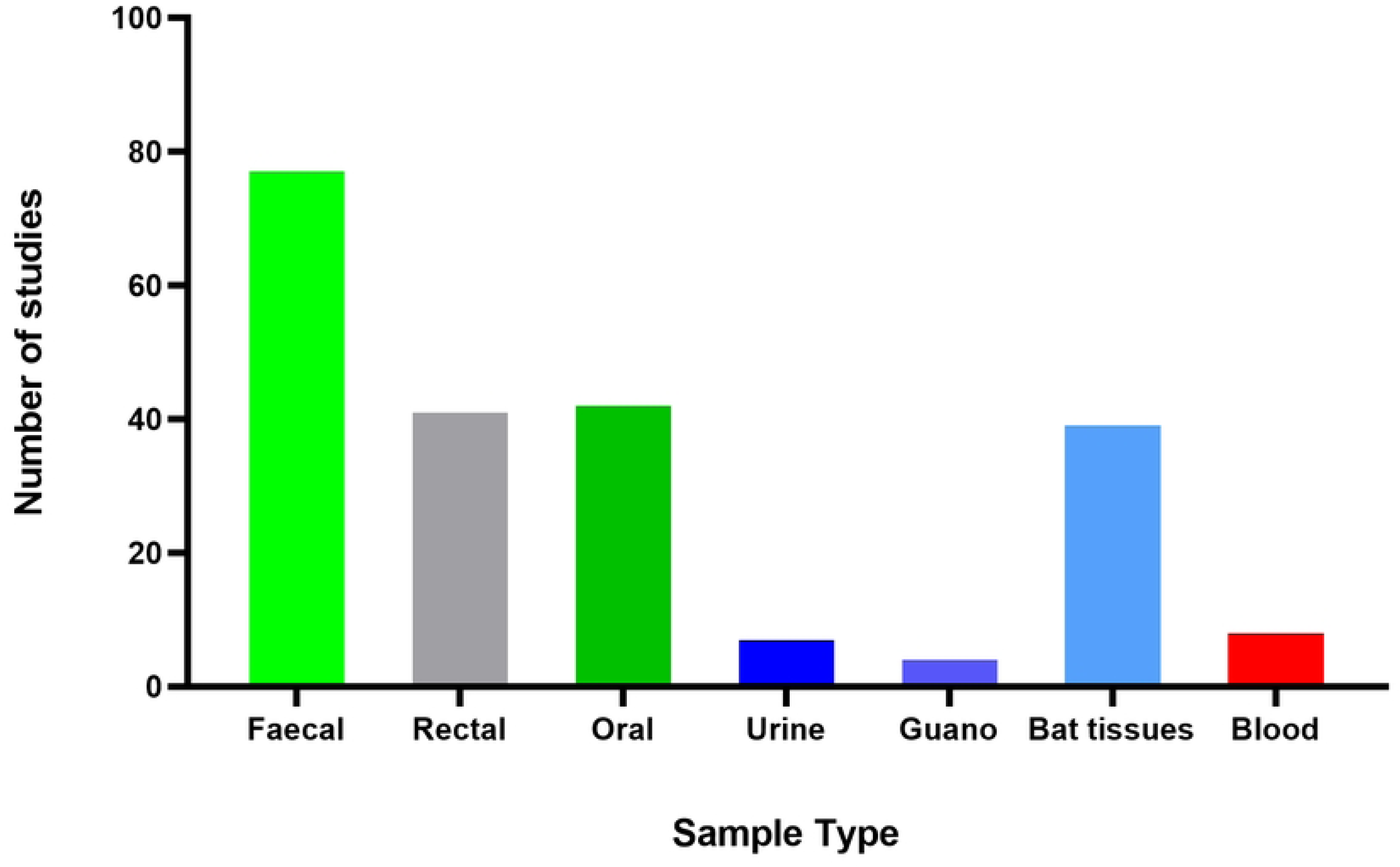
Sample type employed in the bat coronavirus surveillance

**Table 1:**
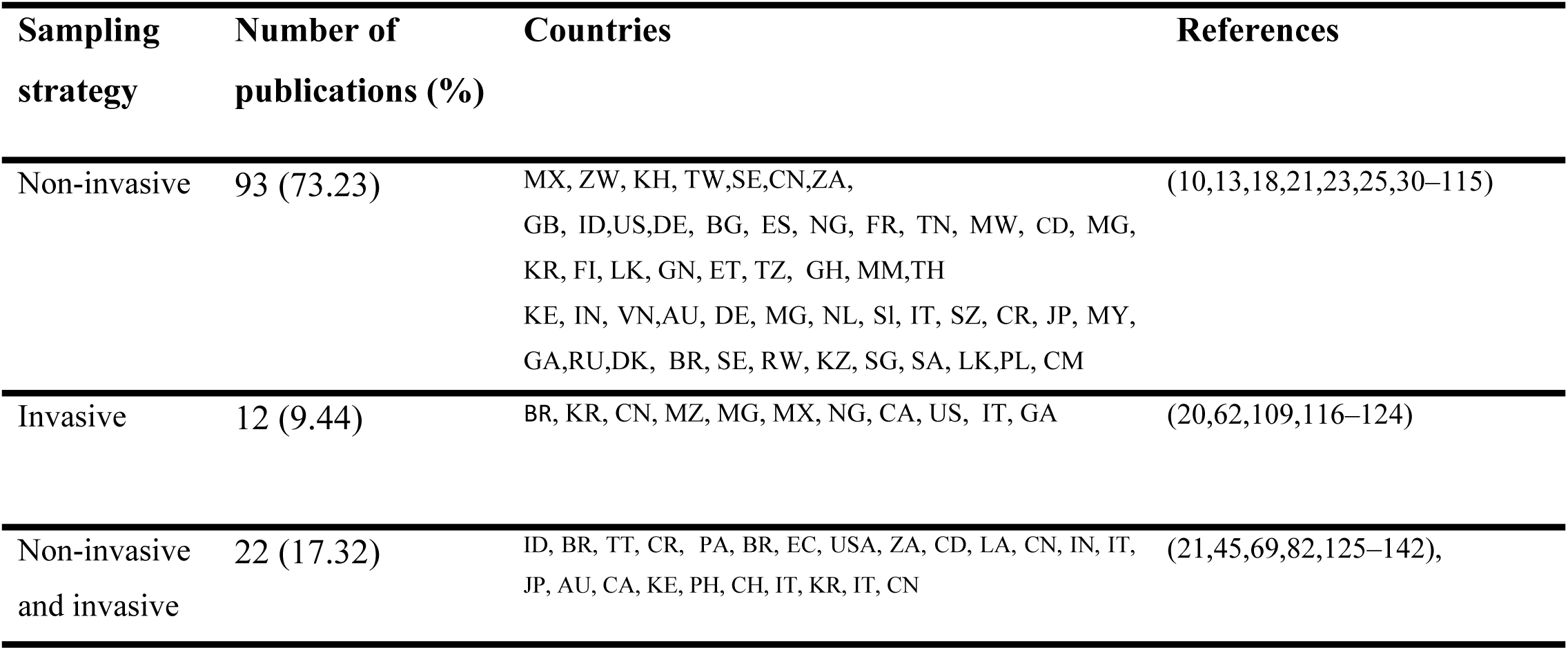
Sampling strategy and sample type employed in the bat coronavirus surveillance.

| Sampling strategy | Number of publications (%) | Countries | References |
| --- | --- | --- | --- |
| Non-invasive | 93 (73.23) | MX, ZW, KH, TW, SE, CN, ZA, GB, ID, US, DE, BG, ES, NG, FR, TN, MW, CD, MG, KR, FI, LK, GN, ET, TZ, GH, MM, TH, KE, IN, VN, AU, DE, MG, NL, SI, IT, SZ, CR, JP, MY, GA, RU, DK, BR, SE, RW, KZ, SG, SA, LK, PL, CM | (10,13,18,21,23,25,30–115) |
| Invasive | 12 (9.44) | BR, KR, CN, MZ, MG, MX, NG, CA, US, IT, GA | (20,62,109,116–124) |
| Non-invasive and invasive | 22 (17.32) | ID, BR, TT, CR, PA, BR, EC, USA, ZA, CD, LA, CN, IN, IT, JP, AU, CA, KE, PH, CH, IT, KR, IT, CN | (21,45,69,82,125–142), |

### 3.3 Bats Species Sampled for Bat Coronavirus Surveillance

Collectively, 324 bat species were sampled from 17 distinct bat families (Table 2). These were *Rhinolophidae* (Horseshoe Bats), *Hipposideridae* (Old World Leaf-nosed Bats), *Phyllostomidae* (Leaf-nosed Bats), *Rhinopomatidae* (Mouse-tailed Bats), *Vespertilionidae* (Vesper Bats), *Emballonuridae* (Sheath-tailed Bats), *Pteropodidae* (Fruit Bats), *Molossidae* (Free-tailed Bats), *Noctilionidae* (Bulldog Bats), *Nycteridae* (Slit-faced Bats), *Mormoopidae* (Ghost-faced Bats), *Megadermatidae* (False Vampire Bats), *Miniopteridae* (Long-winged Bats), *Nandinidae* (Palm Civets), *Natalidae* (Funnel-eared Bats), *Craseonycteridae* (Bumblebee Bat) and *Myonycteridae*.

**Table 2:**
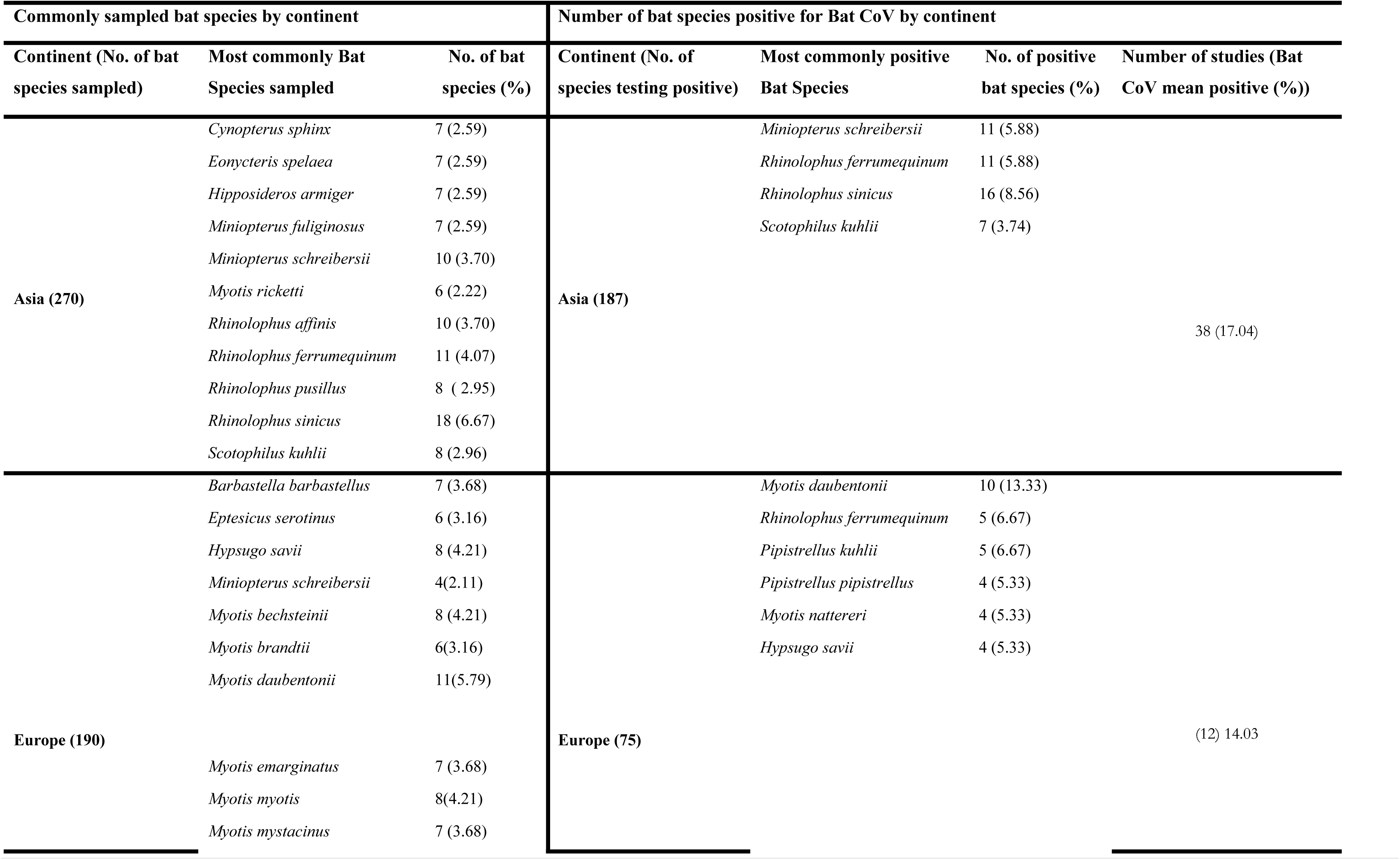
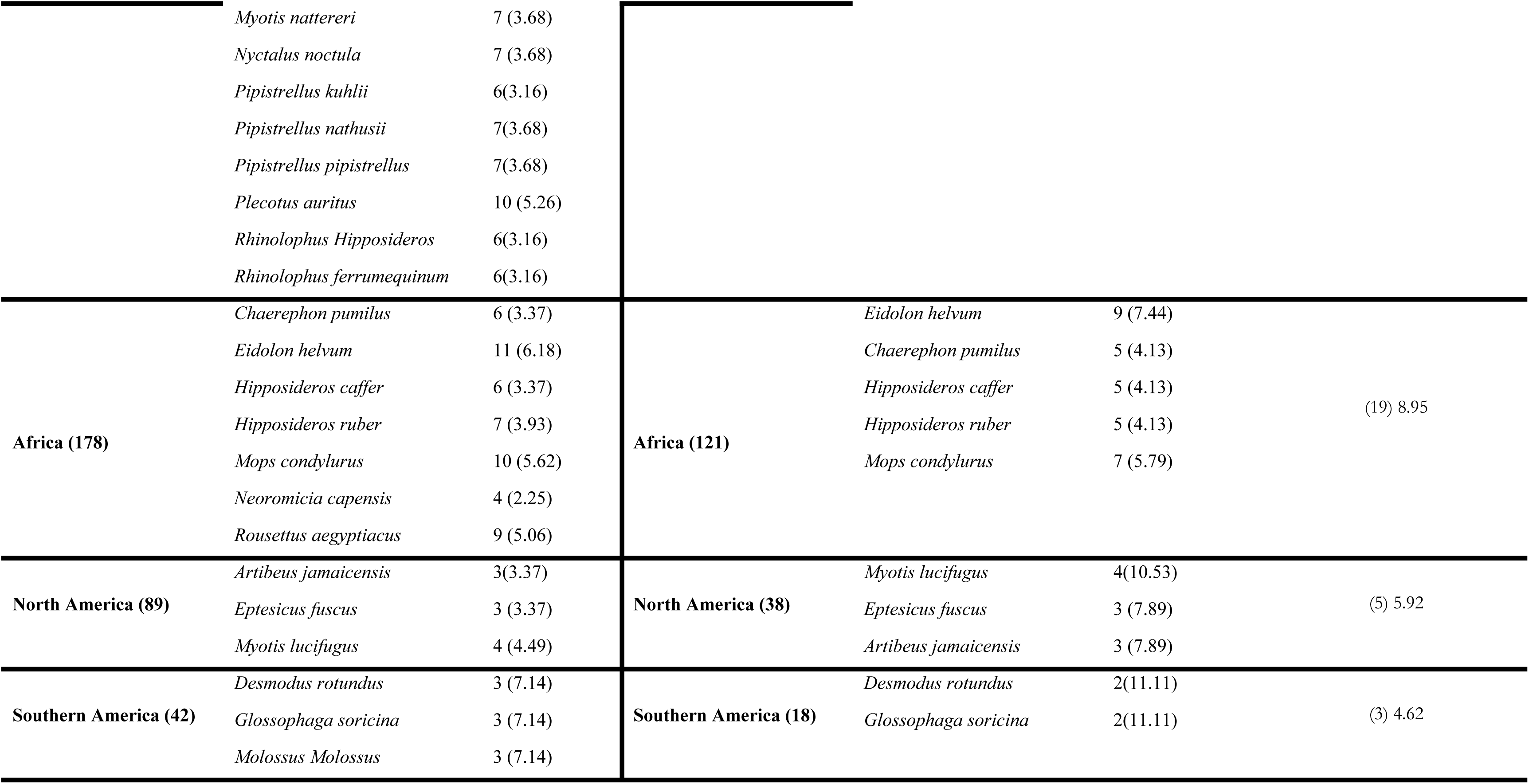
Overview of Bat Species in Studies - Most Commonly Sampled and most commonly bat species testing Positive for Bat CoV.

The bat species with the widest global representation among those studied included Myotis spp., *Rhinolophus sinicus* (commonly sampled in Asia), *Myotis daubentonii* (frequently sampled in Europe), *Eidolon helvum* (predominantly sampled in Africa), *Myotis lucifugus* (commonly sampled in North America), and *Desmodus rotundus* and *Molossus molossus* (frequently sampled in South America).

Two hundred and six (206) bat species tested positive for coronavirus of which 78 were from Asia, 64 from Africa, 29 from Europe, 25 from North America, 13 from South America, and 22 from Australia. The bat species that exhibited predominantly positive occurrences in Asia and Europe were *Rhinolophus sinicus* (Table 2). In Africa, the most prevalent positive species were *Eidolon helvum*, while in North America, it was *Myotis lucifugus*. In South America, the notable species were *Carollia perspercillata* (Table 2)

### 3.4 Molecular detection methods used in bat coronavirus screening and detection

Molecular assays have been used to characterize the bat CoVs and are critical for detecting viruses and predicting their propensity potential in humans. Generally, 94.5% (n = 120/127) of studies used PCR assays that amplified the partial RdRp gene, for detection, yielding partial sequences ranging from 123 to 440 bp. Other targets were helicase 0.8% (n= 1/127), Spike gene(S) 7% (n=9/127), Nucleocapsid gene (N) 2.4% (n=3/127), Envelope gene (E) 2.4% (n= 3/127), and Membrane gene (M) 0.8% (n= 1/127). Full genome sequencing was used in 33.9% of studies (n = 43/127) while metagenomics approach was used in 15.7 % (n= 20/127) studies. Additionally, some of these studies that employed full genome sequencing used the RdRp gene for initial detection. About 55.9% (71) of the studies employed primers that were specific to the target sequence, sometimes known as consensus primers. In contrast, 5.5% (7) of the studies utilized sequence-independent single primer amplification. The remaining 38.6% (49) of studies did not provide information regarding the type of primers used.

## 4.0 DISCUSSION

The prevalence of human emerging infectious diseases (EIDs) is attributed to wildlife, accounting for an estimated range of 60% to 80% [136,145,146]. Extensive scientific investigations have established that bats serve as natural reservoirs of CoVs, which are responsible for respiratory diseases in humans [147–151]. The emergence of zoonotic diseases originating from bats presents a significant peril to global public health, and the repercussions of the pandemic have had a devastating impact on the economies of nations. The COVID-19 pandemic is widely recognized as a significant zoonotic disease of the twenty-first century. It has had a profound impact on global economies, both large and small, due to the implementation of various measures aimed at mitigating and controlling the spread of the virus (150).

It is noteworthy that China is at the forefront of bat CoVs surveillance contributing to over a quarter of the total publications reported across 54 different nations for several reasons. Firstly, China has a rich diversity of bat species, many of which are known to harbour CoVs (151). The country’s diverse biodiversity and different ecosystems create a special habitat for the natural evolution and spread of these viruses. Secondly, China has encountered previous outbreaks of two highly pathogenic human CoVs, SARS-CoV (152) and SARS-CoV-2 (153) as well as other zoonotic viruses [134, [156–158]. This has heightened awareness and prompted proactive measures to monitor and study bat populations as potential reservoirs of CoVs. Additionally, China has invested significantly in scientific research and infrastructure, enabling the establishment of robust surveillance programs.

The surveillance of CoVs in bats commenced subsequent to the occurrence of the initial human CoV pandemic in 2003 (157). Further investigations were undertaken following the emergence of the second highly pathogenic human coronavirus known as MERS-CoV in 2012 [9, 10, 159]. This observation is apparent given that 86.6 % of the studies included in this investigation were published between 2012 and 2022. The observed concentration of studies during this specific timeframe might reflect a heightened global awareness of the potential zoonotic threats posed by CoVs originating from bats. Furthermore, the years 2012 to 2022 were marked by advancements in sequencing technologies and an increased availability of viral genome data.

About three-quarters of the studies used non-invasive sampling. One of the primary advantages of non-invasive methods is their alignment with ethical considerations [160–162]. These methods allow researchers to obtain valuable data while minimizing harm to bat populations and their habitats. This is particularly crucial when studying species that are already vulnerable or endangered. Moreover, non-invasive methods facilitate repeated sampling from the same bat over time. Invasive methods are typically employed when the research necessitates the analysis of internal tissues, organs, or fluids that cannot be obtained non-destructively. However, it comes with ethical and ecological considerations. The removal or harm to bats or their roosting sites must be carefully managed to minimize potential negative impacts. Researchers employing invasive methods should be aware of these ethical considerations and make efforts to ensure their studies contribute to both scientific knowledge and the conservation of bat populations (163).

Faecal samples were the most commonly utilized specimens. The faecal samples were used in 61.4 % of studies examined. The utilization of faecal samples presents numerous benefits for researchers engaged in the study of bat CoVs detection. The collection of faecal samples from bats is a noninvasive procedure that does not inflict harm upon the animals. Moreover, the analysis of faecal samples frequently provides valuable information regarding the presence of viruses and their genetic diversity, given that CoVs are shed in bat faeces. The fact that the virus is successfully isolated from faeces [21, 27, 120, 164] indicates that virus transmission can occur through the faecal route.

Collectively, sampled bat species were drawn from the *Phyllostomidae, Vespertilionidae, Molossidae, Megadermatidae, Hipposideridae, Pteropodidae, Hipposideridae, Nycteridae, Rhinolophidae, Emballonuridae, Craseonycteridae, Noctilionidae, Nycteridae, Mormoopidae, Miniopteridae, Nandinidae*, and *Natalidae* families. These families represented a rich tapestry of bat biodiversity. Notably, *Myotis* species, commonly known as Vesper bats, emerged as the most frequently sampled bat species globally. This finding is consistent with studies that suggest that *Myotis* species are widely distributed (165) and are known reservoirs for various CoV species. Hence, they are important subjects for further research into the dynamics of CoV transmission.

In Asia, *Rhinolophus sinicus* commonly referred to as Horseshoe bats stood out as the species most frequently sampled. The Rhinolophus bats, have been implicated in the transmission of CoVs and are recognized reservoir hosts(166). *Eidolon helvum*, also known as the African Straw-colored Fruit Bat, dominated as the commonly sampled bat species in Africa. These fruit bats are known reservoirs for certain CoVs (167), emphasizing the importance of monitoring their CoV diversity. Africa and other continents are home to a variety of bat species, the fact that the majority of studies are from Asia highlights the inadequate bat CoV surveillance on these continents.

Our findings demonstrate notable regional variations in the positivity rates of bat CoVs. Asia exhibited the greatest prevalence of bat species testing positive for the CoV, accounting for 29.9% of the total studies.

The high positive rate in Asia is consistent with previous research studies that pinpointed Asia as a hotspot for CoV pandemics and carried out rigorous post-pandemic surveillance. Furthermore, there are many different species of bats in the area. The spread of bat CoV can also be attributed to cultural behaviours prevalent in the region, such as the trade of live animals in wet market places [166, 167]. As a consequence, this interaction between animals and humans enhances the likelihood of interspecies transmission of CoV. The presence of coronavirus-positive bat species in the rest of the continent, further emphasizes the global distribution of CoVs among bat populations. Compared to Asia, the recorded lower positivity rates in other continents may be attributed to differences in bat species composition, ecological factors, or fewer surveillance efforts. It is imperative to acknowledge that the lower levels of positivity observed in these areas do not necessarily indicate a lower public health risk, as the existence of a lower prevalence does not eliminate the possibility of zoonotic spillover events.

Molecular techniques have significantly contributed to the identification and characterization of bat CoVs, thereby offering valuable insights into their genetic diversity and evolutionary patterns. About 55.9 % of studies employed target sequence-dependent primers. These primers were designed to be either species-specific, genus-specific, or broad-range [31, 46, 47, 118, 168]. Despite the fact that these assays focus on a conserved region, the current primers may exhibit reduced sensitivity in detecting a wider range of viruses due to the significant genetic diversity observed among CoVs. A scant 5.5% of research papers utilized Sequence-Independent, Single-Primer amplification (SISPA). The SISPA method is a highly effective approach for identifying nucleic acid material, as it does not rely on the precise sequence being amplified or targeted. The PCR assays utilized in the majority of the studies involved the amplification of the RNA-dependent RNA polymerase (RdRp) fragment using primers developed by various researchers [17, 72, 168, 169]. The partial sequences of RdRp pose a challenge in achieving a thorough genomic characterization of viruses at the species level, as it is imperative to observe a threshold value of over 90% amino acid sequence identity in the conserved replicase domain, ADP-ribose-1-phosphatase (ADRP), 3-chymotrypsin like protease (3CLpro), nsp12 (RdRp), helicase, nsp14 exonuclease (ExoN), nsp15 endoribonuclease (NendoU), and nsp16 methyltransferase (O-MT) in order to classify CoVs as members of the same species, thereby contributing to the limited understanding of viral genetic diversity (173). In the context of the analysed publication, it was observed that sequences examined were attributed to either an *alphacoronavirus* or a *betacoronavirus*, based on the availability of complete genome sequence data. The partial RdRp sequences were classified through phylogenetic analysis as closely associated with *alphacoronaviruses* or closely associated with *betacoronaviruses*.

One potential approach to incorporate the partial genomic sequences, as suggested by Auther et al, (18), is to utilize RdRp-grouping units (RGU). This technique enables the elongation of incomplete nucleotide sequences up to a length of 816 nucleotides thereby facilitating the comprehensive analysis of bat coronavirus. Out of the studies that were analysed a number of investigations utilised RGU methodologies, as evidenced by [24, 50, 108], 130] The RGU defines a 4.8% amino acid distance in the analysed 816-bp RdRp fragment for *alphacoronaviruses* and a 6.3% amino acid distance for *betacoronaviruses* (18). The lack of virus isolates obtained directly from bats challenges the complete genomic characterization of these large and highly variable RNA viruses (174). Complete genome sequencing enables the comparison of several genomic features that have been functionally characterized and this can facilitate the virus classification at the species level.

The advent of next-generation sequencing (NGS) technology has enabled the implementation of metagenomics analyses, which allow for the concurrent screening of virome or individual or multiple organisms [40, 43, 74, 88, 172, 173]. Metagenomics methodologies were employed in approximately 15.7% of the studies. The method allows for the rapid detection of viruses belonging to one or multiple viral families in individual or pooled samples [40, 43], 74, 172, 173]. This strategy is crucial for demonstrating that bats harbour a wide variety of zoonotic viruses that have a significant negative impact on both livestock and human health.

Several next-generation sequencing (NGS) technologies have been employed for the detection of bat coronaviruses. These include pyrosequencing [134] sequencing by synthesis (Solexa) [23, 40, 51, 58, 74, 174], Ion Torrent, and the third-generation sequencing method known as Nanopore (82). However, it is important to note that NGS requires complex bioinformatics pipelines, and substantial data storage capabilities [175–177].

However, NGS has the capability to concurrently sequence a vast number of DNA fragments, in contrast to the conventional sequencing technique known as the Sanger method, which is limited to sequencing a single DNA fragment at a time. NGS offers several advantages compared to the conventional Sanger sequencing method. These advantages include a faster turnaround time, enhanced sensitivity, a lower limit of detection, and a more cost-effective approach (174).

## 5.0 Conclusion

The majority of the studies were conducted in Asia, with China exhibiting a greater share in comparison. Molecular approaches used included PCR, coupled with sequencing using Sanger and next-generation technologies and phylogenetic analysis. The majority of studies on bat CoVs utilised PCR assays that targeted the RdRp fragment for virus detection. While effective for initial identification, the use of partial RdRp sequences presents limitations in achieving comprehensive genomic characterisation at the species level. These PCR assays employed primers that were species-specific, genus-specific, or broad-range, each contributing differently to the identification and characterisation of bat CoVs. The virus detection was coupled with sequencing using Sanger or next-generation technologies. Despite the rich diversity of bat species in Africa, Europe, Australasia, Southern America, and North America, there is a notable lack of extensive genomic surveillance for bat CoVs in the region. This research gap leaves much unknown about the potential reservoirs and transmission dynamics of these viruses. The studies reviewed predominantly employed non-invasive sampling methods, with faecal samples being the most frequently collected. Most studies used non-invasive sampling methods and found the viruses highly detectable in faeces, rectal or anal swabs, and intestine samples. While significant progress has been made in the detection and preliminary characterisation of bat coronaviruses, there remains a need for more comprehensive genomic surveillance, particularly in regions with high bat diversity such as Africa, Europe, Australasia, southern America, and North America. Continuous surveillance efforts, coupled with more detailed genomic analyses, are essential for a deeper understanding of the diversity and evolutionary patterns of CoVs, but also evaluate the potential zoonotic transmission of bat CoVs.

## Supporting information

S1 Checklist. PRISMA 2020 checklist.

## Acknowledgements

We express our deepest gratitude to the Department of Para-clinical Studies at the School of Veterinary Medicine, University of Zambia, for their unwavering support and invaluable assistance throughout the implementation of this study.

## Authors’ Contributions

**Conceptualization**: John Amos Mulemena, Katendi Changula.

**Data curation**: John Amos Mulemena, Katendi Changula.

**Formal analysis**: John Amos Mulemena, Katendi Changula, Sichamba Patson.

**Investigation**: John Amos Mulemena.

**Methodology**: John Amos Mulemena, Katendi Changula.

**Supervision**: Katendi Changula, Edgar Simulundu, Walter Muleya.

**Writing—original draft**: John Amos Mulemena.

**Writing—review & editing**: John Amos Mulemena, Katendi Changula, Edgar Simulundu, Benjamin Mubemba, Simbarashe Chitanga

## Conflicts of Interest

The authors declare no conflicts of interest regarding the publication of this paper.

## Notes

### Competing Interest Statement

The authors have declared no competing interest.

